# A genetic screen to uncover molecular mechanisms underlying lipid transfer protein function at membrane contact sites and neurodegeneration

**DOI:** 10.1101/2023.07.28.550979

**Authors:** Shirish Mishra, Vaishnavi Manohar, Shabnam Chandel, Tejaswini Manoj, Subhodeep Bhattacharya, Nidhi Hegde, Vaisaly R Nath, Harini Krishnan, Corinne Wendling, Thomas Di Mattia, Arthur Martinet, Prasanth Chimata, Fabien Alpy, Padinjat Raghu

## Abstract

Lipid transfer proteins mediate the transfer of lipids between organelle membranes in eukaryotes and loss of function in these has been linked to neurodegenerative disorders. However, the mechanism by which loss of lipid transfer protein function leads to neurodegeneration is not understood. In *Drosophila* photoreceptors, depletion of Retinal Degeneration B (RDGB), a phosphatidylinositol transfer protein localized to endoplasmic reticulum-plasma membrane contact sites leads to defective phototransduction and retinal degeneration but the mechanism by which RDGB function is regulated and the process by which loss of this activity leads to retinal degeneration is not understood. RDGB is localized to membrane contact sites (MCS) and this depends in the interaction of its FFAT motif with the ER integral protein VAP. To identify regulators of RDGB function *in vivo*, we depleted more than 300 VAP interacting proteins and identified a set of 52 suppressors of *rdgB*. The molecular identity of these suppressors indicates a role for novel lipids in regulating RDGB function and for transcriptional and ubiquitination processes in mediating retinal degeneration in *rdgB*. The human homologs of several of these molecules have been implicated in neurodevelopmental diseases underscoring the importance of VAP mediated processes in these disorders.

## Introduction

The maintenance of exact membrane lipid composition is important for providing distinct identity to cellular organelles and thus support normal cellular physiology(Harayama and Riezman, 2018). Various lipid species reach their specific organelle membrane either via vesicular or non-vesicular transport. Proteins that shuttle lipids in a non-vesicular manner across various compartments are known as lipid transfer proteins (LTPs). Each of these LTPs transfer specific lipid species such as sterols, ceramides or phospholipids and in many cases the LTPs are localized at very specific locations known as membrane contact sites (MCS). In a eukaryotic cell, MCS are regions where two organelle membranes come very close at the range of 10-30 nm but do not fuse (Prinz et al., 2020). Being the largest cellular organelle, the endoplasmic reticulum (ER) forms MCS with the mitochondria, lysosomes, Golgi network, lipid droplets and the plasma membrane (PM). MCS provide fast and efficient delivery of metabolites between two membranes and could be permanent or induced (Wu et al., 2018); this includes the exchange of lipids between organelle membranes to support ongoing cell physiology (Cockcroft and Raghu, 2018). Growing evidence suggest an important role for LTP function at MCS and LTPs in human neurological disorders (Peretti et al., 2020) (Fowler et al., 2019) (Guillén-Samander and de Camilli, 2022). However, much remains to be discovered on the regulation of LTP function at MCS.

MCS between the ER and the PM are important for regulating both plasma membrane lipid composition and signalling functions. One of the best examples for the requirement of an LTP at the ER-PM MCS is sensory transduction in *Drosophila* photoreceptors (Yadav et al., 2016). Photoreceptors detect light through the G-protein coupled receptor (GPCR) rhodopsin (Rh), leading to the hydrolysis of phosphatidylinositol 4,5-bisphosphate [PI(4,5)P_2_] by G-protein coupled phospholipase C (PLC) activity (Hardie and Raghu, 2001). As part of their ecology, fly photoreceptors are exposed to light; in bright daylight they typically absorb ca. 10^6^ effective photons/second resulting in extremely high PLC activity. Hence, fly photoreceptors provide an excellent model system to study the turnover of PI(4,5)P_2_ during PLC mediated cell signalling (Raghu et al., 2012).

Given the low abundance of PI(4,5)P_2_, replenishment of this lipid at the PM is necessary for uninterrupted PLC signalling. Many enzymes and proteins participate in this process but a key step is the transfer of lipids that are intermediates of the PI(4,5)P_2_ cycle. One of the proteins at this site is <u>R</u>etinal <u>D</u>egeneration <u>B</u> (RDGB), a large multi-domain protein with an N-terminal phosphatidylinositol transfer protein (PITP) domain (Raghu et al., 2021). The PITP domain belongs to the superfamily of LTPs. In the case of RDGB, its PITP domain can transfer phosphatidylinositol (PI) and phosphatidic acid (PA) *in vitro* (Yadav et al., 2015a) a property that is conserved in its mammalian ortholog, Nir2 (Kim et al., 2015)13. *rdgB* mutant flies undergo light dependent retinal degeneration, a reduced ERG response and a reduced rate of PI(4,5)P_2_ resynthesis at the PM following PLC activation (Harris and Stark, 1977; Hotta and Benzer, 1970; Yadav et al., 2015b)In photoreceptors, RDGB is localized at the ER-PM MCS formed between the microvillar plasma membrane and the sub-microvillar cisternae (SMC), a specialization of the smooth endoplasmic reticulum (Yadav et al., 2016)8. The localization of RDGB at this MCS is critically dependent on its interaction with the ER integral membrane protein VAP. This interaction is physiologically relevant as disruption of the protein-protein interaction between RDGB and VAP in *Drosophila* photoreceptors results in mislocalization of RDGB from this MCS, reduced the efficiency of PI(4,5)P_2_ turnover and impacts the response to light (Yadav et al., 2018)16. However, the mechanisms by which the activity of RDGB is regulated by other proteins at the MCS in this *in vivo* model system remains to be discovered. VAP proteins are involved in a range of interactions with proteins containing FFAT/FFNT/Phospho-FFAT/non-FFAT motifs (Cabukusta et al., 2020; di Mattia et al., 2020; Slee and Levine, 2019). Thus, it seems possible that other proteins involved in regulating biochemical activity at this MCS might also be localized to the SMC via VAP interactions. The identification and analysis of proteins engaged in VAP dependent interactions might help in understanding the regulation of RDGB function. Importantly, VAP proteins have been implicated in neurodegenerative disorders such as amyotrophic lateral sclerosis (ALS), Frontotemporal dementia (FTD), Alzheimer’s disease (AD) and Parkinson’s disease [reviewed in (Dudás et al., 2021)].

In this study, we have carried out a proteomics screen to identify protein interactors of VAP-A and VAP-B in mammalian cells and tested their function significance in the context of neurodegeneration using the experimental paradigm of RDGB function in *Drosophila* photoreceptors *in vivo*. The candidates so identified perform a wide range of sub-cellular functions indicating an extensive network of biochemical processes that control the function of RDGB in regulating lipid transfer during PLC signalling, thus maintaining the structural and functional integrity of neurons.

## Results

### Strategy of Proteomics screen

To obtain a list of proteins interacting with VAPs, we performed pull-down experiments in human cells. We produced, in *Escherichia coli*, and purified the MSP domain of human VAP-A and VAP-B fused to C-terminal 6His tag (Fig 1A). As negative control, we used the K94D/M96D and K87D/M89D mutants (herein named KD/MD mutants) of VAP-A and VAP-B, respectively, that are unable to bind FFAT (two phenylalanine in an acidic tract) motifs ((Kaiser et al., 2005; Wilhelm et al., 2017)21,22). Each recombinant protein was attached to a Ni^2+^-NTA resin, and then incubated with protein extracts from HeLa cells. Bound proteins were eluted and analyzed by SDS–PAGE followed by silver nitrate staining (Fig 1B) that showed numerous differential bands between wild type (WT) and mutant VAP samples, suggesting that many proteins are pulled down owing to VAP’s ability to bind FFAT motifs. To verify the pull-down efficiency, we performed Western blot using antibodies against two known VAP partners, ORP1 and STARD3NL (Fig 1C) (Alpy et al., 2013; Rocha et al., 2009). ORP1 exists as a long and a short isoform called ORP1L and ORP1S respectively, ORP1L being the only one of the two to possess a FFAT motif. As expected, the ORP1L isoform was pulled down by WT VAPs but not by mutant VAPs, and the ORP1S isoform was not precipitated (Fig. 1C). Besides, STARD3NL co-precipitated with WT VAP-A and VAP-B and not with mutant VAPs, while actin, used as a loading control, was not found in the eluted fractions (Fig. 1C). To identify the proteins pulled down by VAPs, elutions were analyzed by tandem mass spectrometry (MS/MS). To identify proteins pulled down according to their ability to interact with VAPs in an FFAT-dependent manner, proteins were ranked based on their enrichment in the WT over the KD/MD mutant VAP sample, and on their MS/MS score (Fig. 1D). This strategy led to the identification of 403 proteins, 193 of which were pulled-down by both VAP-A and VAP-B. Interestingly, many known partners of VAP-A and VAP-B, such as OSBP, ORP1, ORP2, WDR44, VPS13A, VPS13D were identified (Fig. 1D). Using a position weight matrix strategy, we looked for potential FFAT and Phospho-FFAT in the protein sequences; sequences were attributed a score, with 0 corresponding to an ideal FFAT/Phospho-FFAT sequence. Among the 403 proteins identified, 136 had a FFAT or Phospho-FFAT with a significant score (between 0 and 2.5) (Sup Table S1). We used this list of 403 mammalian proteins and identified their *Drosophila* orthologs using the DRSC integrative ortholog prediction tool (DIOPT) (Hu et al., 2011) and the fly orthologs with the best score were identified. Using this approach, we were able to identify Fly orthologs with more than 90% coverage for 393 out of 403 mammalian proteins in the VAP interaction list (Sup. Table S2).

**Figure 1:**
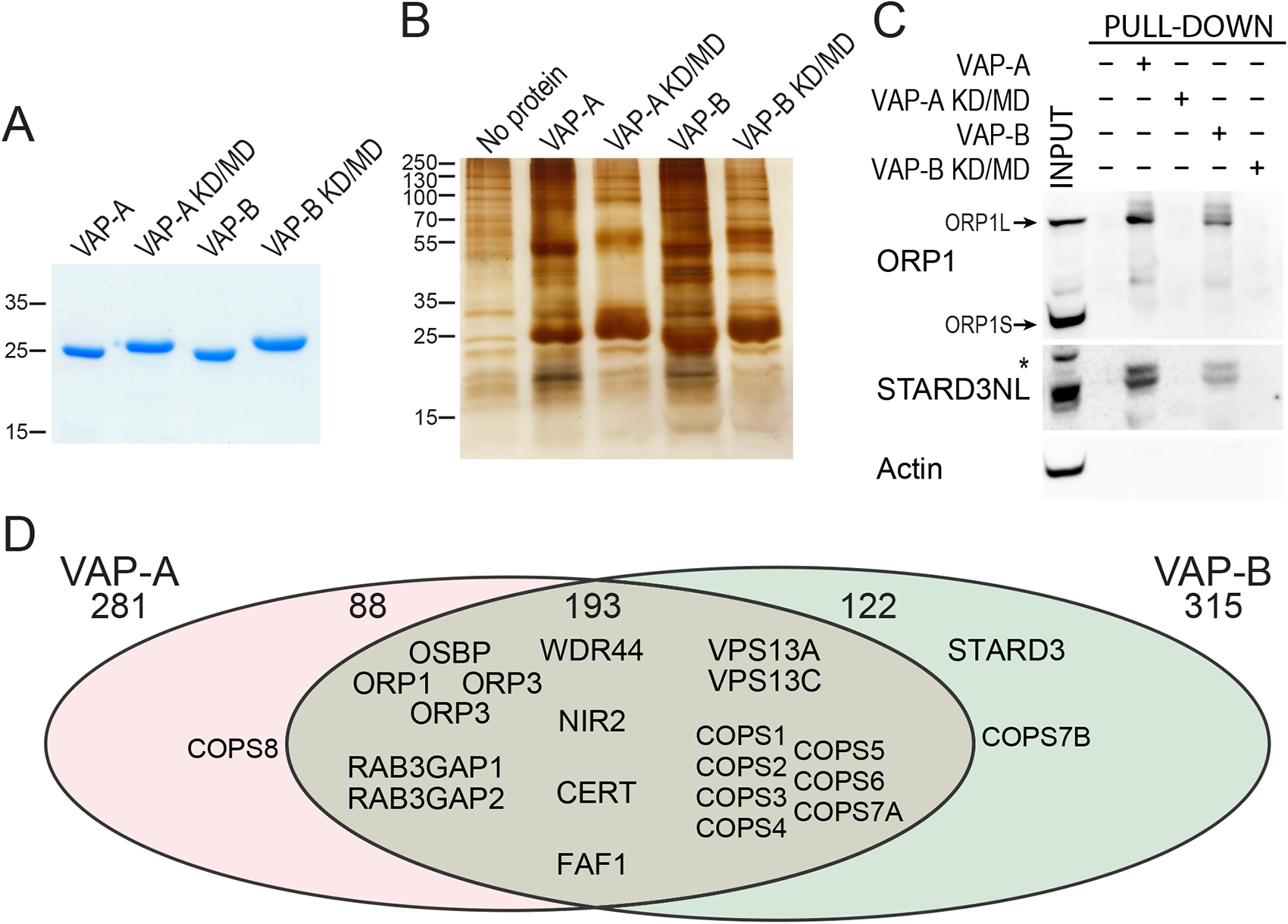
Identification of VAP-A and VAP-B binding partners. (A) Coomassie Blue staining of the recombinant WT and KD/MD mutant MSP domains of VAP-A and VAP-B after SDS–PAGE. (B) Silver nitrate staining of proteins pulled down using WT MSP domains of VAP-A and VAP-B, and the KD/MD mutant MSP domains, after SDS–PAGE. (C) Western blot analysis of proteins pulled down using the WT and mutant MSP domain of VAP-A and VAP-B. The input and pull-down fractions correspond to HeLa cell total protein extract and bound proteins, respectively. *: non-specific band. D: Venn diagram of proteins pulled-down by VAP-A and VAP-B (and not by mutant VAP-A and VAP-B). A total of 403 proteins were pulled-down with either VAP-A or VAP-B. 193 proteins were pulled-down with both VAP-A and VAP-B.

### Strategy of genetic screen

To identify *in vivo* regulators of RDGB function at ER-PM contact sites, we utilized a hypomorphic allele *rdgB^9^* (Vihtelic et al., 1991). *rdgB^9^* expresses a small amount of residual RDGB protein that provides some function in contrast to the protein null allele *rdgB^2^*. The FFAT motif of RDGB interacts with the ER resident membrane protein dVAP-A to provide both localization and function to RDGB (Yadav et al., 2018). FFAT motifs are found in many proteins of varied biological functions and serve to localize them to ER contact sites through a protein-protein interaction with VAP (Murphy and Levine, 2016). We reasoned that if several proteins with an FFAT motif bind to VAP at the ER-PM interface, the lipid transfer function of RDGB could be modulated by their presence at the ER-PM MCS (Figure 2A). Such proteins, relevant to RDGB function could be identified by testing their ability to modify the phenotype of the *rdgB* mutant.

**Figure 2:**
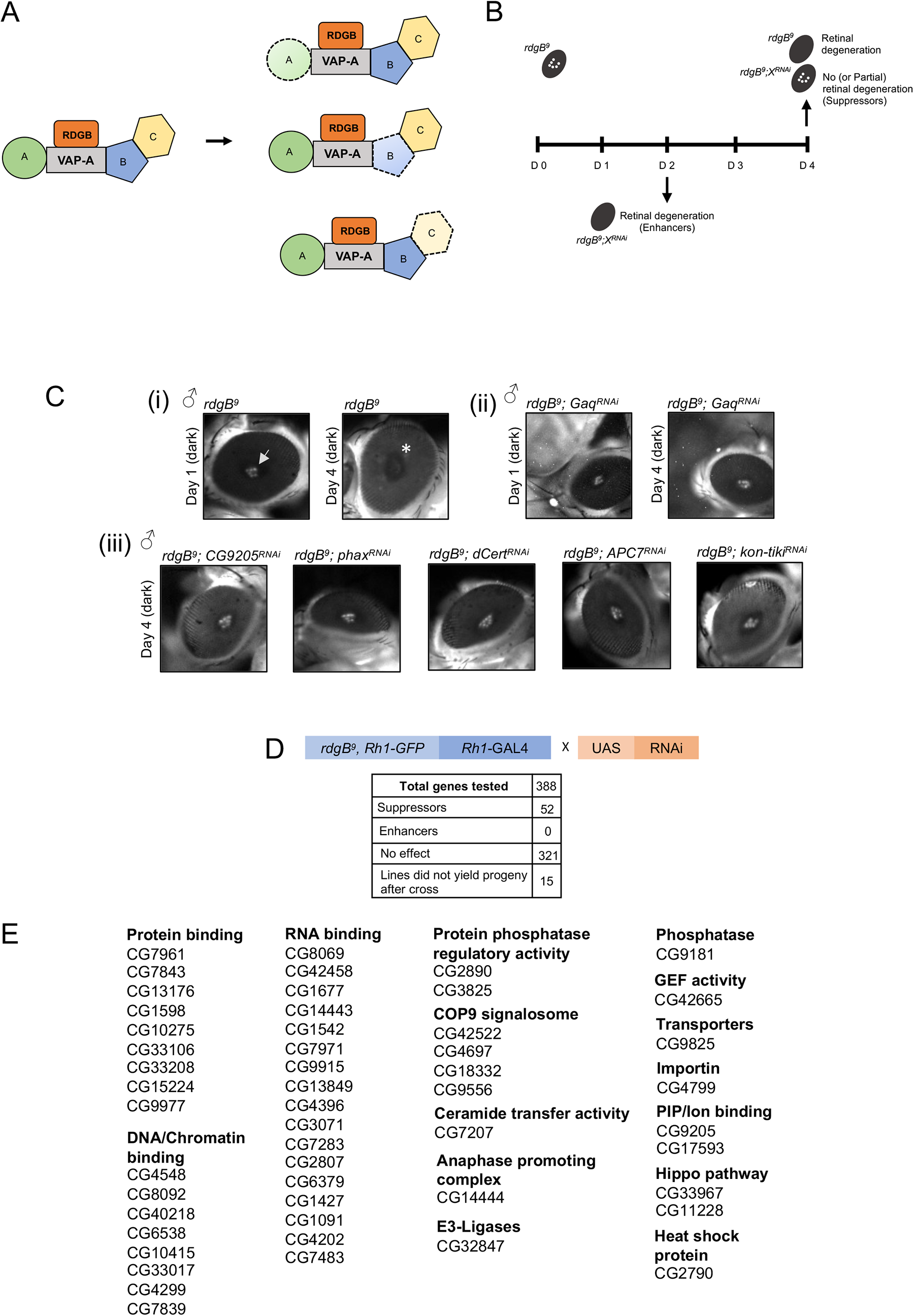
Strategy of the genetic screen and hits found. (A) Cartoon depicting classes VAP interactors used in the present genetic screen. Depletion of a specific VAP interactor is depicted with dotted line. Fly homologues were filtered using DIOPT in Flybase (http://flybase.org/). (B) Genetic scheme used to find either enhancers or suppressors of the retinal degeneration phenotype of *rdgB^9^*. (C) Pseudopupil imaging (i) *rdgB^9^* showed retinal degeneration by day four in dark when checked via deep pseudopupil imaging. (ii) The degeneration was partially suppressed when levels of Gaq were downregulated in *rdgB^9^* on day four. (iii) Selected hits that showed suppression of retinal degeneration in *rdgB^9^* on day four. (D) Table showing the full list of genes used in the screen and number of suppressor genes identified. (E) Positive hits (suppressor genes) are divided in different categories depending on their cellular functions. n=5 flies/RNAi line

*rdgB^9^* shows retinal degeneration that is enhanced when flies are grown under illumination (Harris and Stark, 1977; Stark et al. 1983). Under illumination, *rdgB^9^* flies show severe retinal degeneration by two days post eclosion making it difficult to score for modulation of this phenotype by other gene products. To overcome this problem, we reared *rdgB^9^* flies without illumination, a condition under which the retinal degeneration still occurs but at a slower rate; in dark reared *rdgB^9^* flies it takes two days for the retinal degeneration to set in and by day four complete retinal degeneration was seen (Figure 2B). Retinal degeneration was scored by visualizing the deep pseudopupil (DPP) under a fluorescence stereomicroscope(Georgiev et al., 2005)29. To visualize fluorescent pseudopupil, a protein fusion of Rhodopsin1 (Rh1) was tagged with GFP, expressed under its own promoter, and recombined in *rdgB^9^*. Under these conditions *rdgB^9^* shows a clear fluorescent DPP on day 1 that is lost by day 4 with the progression of retinal degeneration (Figure 2C i).

To identify molecules regulating RDGB function, we depleted their mRNA levels using transgenic RNAi from publicly available collections (Dietzl et al., 2007; Perkins et al., 2015); for 5 out of 393 fly genes there were no RNAi line available from public resources (Sup. Table S2). The eye specific Rh1 promoter was used to restrict GAL4 expression and thus gene depletion, in space to the outer six photoreceptors and in time to post 70 hrs pupal development (Yadav et al., 2015). To validate the genetic screen, *G*_αq_ was downregulated in the *rdgB^9^* flies and the pseudopupil was scored after day 2 and 4. Knocking down *G*_αq_ in *rdgB^9^* flies under Rh1 promoter showed partial suppression of retinal degeneration and hence pseudopupil presence after day 4 in dark suggested the efficacy of the screening method (Figure 2C ii).

Using this strategy, we depleted each of the 388 VAP interacting proteins via RNAi in the *rdgB^9^* sensitized background (Figure 2C iii, Sup. Table S2). The screen was performed such that the phenotype arising from off targets could be minimized. We first used a single RNAi line per gene of interest for the pseudopupil analysis and once a positive phenotype was scored, the assay was repeated with a second independent RNAi line for the same gene. Only those genes were finally tabulated where two independent lines per gene showed a positive phenotype. To assay the enhancement of retinal degeneration, fly eyes were visualized on day 2 while for suppression, fly eyes were checked on day 4. Any suppresser that showed complete recovery of DPP was scored as a full rescue while others were designated as partial suppressers.

Out of 388 genes, knockdown of 52 (two independent RNAi lines per gene) in *rdgB^9^* showed suppression of retinal degeneration (Figure 2D, Table 1); we designated these as *su(rdgB)*. In this study, we did not identify any candidate that showed enhancement of degeneration when depleted in *rdgB^9^*. Moreover, 15 genes where only a single RNAi line was available, when tested, did not result in adult progeny (larval death/ pupae formed but no fly emerged). Based on their Gene Ontology tags, the 52 *su(rdgB)* could be classified into several categories (Figure 2E). Of these, the largest number of suppressers were from the class of RNA binding and DNA/chromatin binding proteins. Examples of candidates with strong suppression phenotypes are pleckstrin-homology (PH)-domain containing protein (CG9205), phosphorylated adaptor for RNA export (PHAX), ceramide transfer protein (Cert), anaphase promoting complex 7 protein (APC7) and laminin G domain containing protein Kon-tiki (Fig 2 Ciii). These findings indicate that the mechanisms underlying retinal degeneration in *rdgB^9^* likely involve diverse sub-cellular processes.

**Table 1:** List of *rdgB* interactors. Summary of *Drosophila rdgB* genetic interactors identified in the screen. Gene name and/or CG number in Flybase (www.flybase.org), Uniport (https://www.uniprot.org/) accession number along with their GO functional annotation. For each gene the I.D of RNAi lines from VDRC or TRiP library used are shown. Phenotypes scored following depletion of each gene are represented under ‘Suppression’ column; ‘++’ denoted definite suppression while ‘+’ denotes partial suppression. Human orthologue of each *rdgB* interactor is identified. Known phenotypes associated with each human homolog is denoted along with the online Mendelian Inheritance in Man (OMIM) identifier number.

| Total genetic interactors | Primary accession number (UniProt) | 1st RNAi line ID | Suppression | 2nd RNAi line ID | Suppression | Function | Human orthologue | Associated phenotypes | OMIM Number |
| --- | --- | --- | --- | --- | --- | --- | --- | --- | --- |
| CG8069 | A1Z7P3 | 100778/KK | ++ | 28189/GD | ++ | Phosphorylated adaptor for RNA export | PHAX |  | 604924 |
| CG4548 | Q9GQN5 | 101568/KK | ++ | 10618/GD | ++ | XNP/ Adenosinetriphosphatase | ATRX | Alpha-thalassemia/mental retardation syndrome | 300032 |
| CG7961 | Q9W0B8 | 35305/GD | ++ | 35306/GD | + | Coat Protein (coatomer) α | COP-A | Autoimmune interstitial lung, joint, and kidney disease | 601924 |
| CG7843 | Q9V9K7 | 106344/KK | + | 22574/GD | + | Arsenic resistance protein 2 | SRRT (Isoform 5) |  | 614469 |
| CG42665 | Q9VVC6 | 105885/KK | + | 101144/KK | + | Ephexin | ARHGEF5 | Breast cancer | 600888 |
| CG8092 | A0A0B4KER0 | 28196/GD | + | TRiP 25971 | + | relative of woc | POGZ (Isoform 5) | White-Sutton syndrome | 614787 |
| CG42458 | Q7KU81 | 106608/KK | ++ | 108072/KK | ++ | UN, mRNA binding | HNRNPC (Isoform 4) |  | 164020 |
| CG42522 | Q7KTH8 | TRiP 33370 | ++ | No 2nd RNAi available |  | COP9 signalosome subunit 8 | COPS8 (Isoform 2) |  | 616011 |
| CG1677 | Q9W3R9 | 109697/KK | ++ | 50195/GD | + | UN, Predicted to be involved in mRNA splicing, via spliceosome | ZC3H18 (Isoform 2) |  | Not applicable |
| CG14443 | Q9W3Y5 | 105254/KK | ++ | 17618/GD | ++ | UN, RNA helicase | DDX21 |  | 606357 |
| CG1542 | Q9V9Z9 | 104575/KK | ++ | 39976/GD | ++ | UN, Predicted to be involved in rRNA processing and ribosomal large subunit biogenesis | EBNA1BP2 |  | 614443 |
| CG9825 | Q9W1Z1 | 105868/KK | ++ | 1712/GD | ++ | UN, Solute carrier family 17 (SLC17) member | SLC17A7 |  | 605208 |
| CG9205 | Q9W0K9 | 107612/KK | + | 29079/GD | ++ | UN, Oxysterol binding protein; PH domain | OSBPL11 |  | 606739 |
| CG7971 | A8JNI2 | 101384/KK | ++ | 34262/GD | + | UN, Predicted to be involved in RNA splicing | SRRM2 | Intellectual developmental disorder, autosomal dominant 72 | 606032 |
| CG4799 | P52295 | 102627/KK | + | 32466/GD | ++ | Pendulin | KPNA6 |  | 610563 |
| CG9915 | A8JV07 | 103731/KK | ++ | No 2nd RNAi available |  | UN, Predicted to be involved in poly(A)+ mRNA export from nucleus. | IWS1 (Isoform 2) |  | Not applicable |
| CG13849 | Q95WY3 | 103738/KK | + | 51775/GD | + | Nop56 | NOP56 | Spinocerebellar ataxia 36 | 614154 |
| CG9181 | Q9W0G1 | 108888/KK | ++ | 37436/GD | ++ | Protein tyrosine phosphatase 61F | PTPN12 | Colon cancer, somatic | 600079 |
| CG4396 | Q9VYI0 | 101508/KK | + | 48891/GD | + | found in neurons | ELAVL1 |  | 603466 |
| CG33967 | Q9VFG8 | 106507/KK | ++ | 100765/KK | ++ | kibra | WWC1 | Memory, enhanced, QTL | 610533 |
| CG13176 | Q7JW27 | 39769/GD | ++ | 24642/GD | ++ | washout | WASH6P |  | Not applicable |
| CG3071 | Q9W4Z9 | 107206/KK | ++ | 29589/GD | + | UN, Predicted to have snoRNA binding activity | UTP15 |  | 616194 |
| CG1598 | Q7JWD3 | 110555/KK | + | 32391/GD | ++ | Unnamed/ Adenosinetriphosphatase | GET3 | Cardiomyopathy, dilated, 2H | 601913 |
| CG40218 | Q8SXI2 | 102960/KK | ++ | No 2nd RNAi available |  | Yeti | CFDP1 |  | 608108 |
| CG4697 | Q9VJR9 | 34308/GD | ++ | 34307/GD | ++ | COP9 signalosome subunit 1a | GPS1 |  | 601934 |
| CG14444 | Q9W3Y6 | 110729/KK | ++ | 17622/GD | ++ | Anaphase Promoting Complex subunit 7 | ANAPC7 (Isoform 2) | Ferguson-Bonni neurodevelopmental syndrome | 606949 |
| CG2890 | Q9W2U4 | 105399/KK | ++ | 25445/GD | ++ | Protein phosphatase 4 regulatory subunit 2-related protein | PPP4R2 (Isoform 3) |  | 613822 |
| CG7283 | Q9VTP4 | 109345/KK | ++ | 23459/GD | ++ | Ribosomal protein L10Ab | RPL10A |  | 615660 |
| CG2807 | Q9VPR5 | 110091/KK | + | 25162/GD | ++ | Splicing factor 3b subunit 1 | SF3B1 | Myelodysplastic syndrome, somatic | 605590 |
| CG6538 | P41900 | 110569/KK | + | 12602/GD | + | Transcription factor TFIIFβ | GTF2F2 |  | 189969 |
| CG18332 | Q8SYG2 | 101516/KK | + | 12821/GD | ++ | COP9 signalosome subunit 3 | COPS3 |  | 604665 |
| CG6379 | Q9W4N2 | 103723/KK | + | 29611/GD | ++ | Unnamed/ Methyltransferase cap1 | CMTR1 |  | 616189 |
| CG1427 | Q9VNE3 | 105727/KK | + | 17456/GD | ++ | Sec synthetase | SEPSECS (Isoform 3) | Pontocerebellar hypoplasia type 2D | 613009 |
| CG10275 | Q9VJ82 | 106680/KK | ++ | 37283/GD | ++ | kon-tiki | CSPG4 |  | 601172 |
| CG2790 | Q9W0X8 | 101619/KK | + | 20903/GD | + | UN, The Heat Shock Protein 40 (Hsp40) family of co-chaperones | DNAJC21 | Bone marrow failure syndrome 3 | 617048 |
| CG10415 | O96880 | 100572/KK | + | 12592/GD | + | Transcription factor IIEα | GTF2E1 |  | 189962 |
| CG11228 | Q8T0S6 | 104169/KK | ++ | 7823/GD | + | Hippo | STK3 |  | 605030 |
| CG1091 | Q9VI58 | 107175/KK | + | 16088/GD | ++ | Tailor, RNA uridylyltransferase | TUT1 |  | 610641 |
| CG33106 | Q9VCA8 | 103411/KK | ++ | 33394/GD | + | mask, multiple ankyrin repeats single KH domain | ANKRD17 (Isoform 6) | Chopra-Amiel-Gordon syndrome | 615929 |
| CG33208 | Q86BA1 | 105837/KK | + | 25371/GD | + | Mical, Molecule interacting with CasL | MICAL3 |  | 608882 |
| CG15224 | P08182 | 106845/KK | + | 32377/GD | + | Casein kinase II β subunit | CSNK2B | Poirier-Bienvenu neurodevelopmental syndrome | 115441 |
| CG17593 | Q9VQR9 | 106469/KK | ++ | 13029/GD | ++ | UN, Orthologous to human CCDC47 (coiled-coil domain containing 47) | CCDC47 | Trichohepatoneurodevelopmental syndrome | 618260 |
| CG33017 | A1ZAC8 | 103968/KK | + | 40022/GD | + | UN, The MADF-BESS domain transcription regulators | GPATCH8 (Isoform 2) |  | 614396 |
| CG4299 | P53997 | 108987/KK | + | TRiP 77433 | + | Set, encodes a subunit of the inhibitor of histone acetyltransferase (INHAT) complex | SET | Intellectual developmental disorder, autosomal dominant 58 | 600960 |
| CG7207 | Q9Y128 | 103563/KK | ++ | 27914/GD | + | ceramide transfer protein | CERT | Intellectual developmental disorder, autosomal dominant 34 | 604677 |
| CG4202 | Q9I7W5 | 103352/KK | ++ | 49946/GD | + | Something about silencing 10 | UTP3 |  | 611614 |
| CG9977 | Q9VZX9 | 106749/KK | ++ | 36193/GD | ++ | Adenosylhomocysteinase like 1 | AHCYL1 |  | 607826 |
| CG32847 | Q8IQM1 | 104294/KK | ++ | 48423/GD | ++ | UN, contains The RING (Really Interesting New Gene) finger domain | TRIM26 |  | 600830 |
| CG7839 | Q9VTE6 | 105979/KK | + | 12691/GD | + | UN, Orthologous to human CEBPZ (CCAAT enhancer binding protein zeta). | CEBPZ |  | 612828 |
| CG7483 | Q9VHS8 | 108580/KK | ++ | TRiP 32444 | + | eIF4AIII, ATP-dependent RNA helicase | EIF4A3 | Robin sequence with cleft mandible and limb anomalies | 608546 |
| CG9556 | Q94899 | 48044/GD | ++ | TRiP 28908 | ++ | alien | COPS2 |  | 604508 |
| CG3825 | Q9W1E4 | 107545/KK | + | TRiP 33011 | + | Protein phosphatase 1 regulatory subunit 15 | PPP1R15B | Microcephaly, short stature, and impaired glucose metabolism 2 | 613257 |

### Identification of suppressors specific to *rdgB^9^*

In principle, depletion of a gene product can suppress retinal degeneration in *rdgB^9^* by one of two mechanisms (i) by altering the underlying biochemical abnormality resulting from loss of RDGB function, i.e. the trigger (ii) by downregulating downstream sub-cellular processes that are part of the degenerative process, i.e the effectors. Genes in the first category, i.e the trigger mechanism, might be expected to suppress only the degeneration of *rdgB^9^* and no other retinal degenerations whereas gene that are effectors of retinal degeneration might be expected to suppress multiple retinal degeneration mutants.

To distinguish these two categories of genes we tested each of the 52 *su(rdgB)* for their ability to block retinal degeneration in *norpA^p24^* (Figure 3A, Sup. Table S3). *norpA* encodes for the PLC and catalyzes the hydrolysis of PI(4,5)P_2_ to DAG and IP_3_. *norpA^p24^* is a strong hypomorph and show light dependent retinal degeneration(Pearn et al., 1996). Out of the 52 *su(rdgB)*, 13 genes partially suppressed light dependent retinal degeneration in *norpA^p24^* suggesting that they likely participate in the process of retinal degeneration (Figure 3B, 3C). Most genes in this category belong to the class of RNA binding/processing and DNA/ Chromatin binding (Figure 3C). The remaining 39 genes therefore likely represent unique suppressors of *rdgB^9^* and therefore may participate specifically in the trigger mechanism.

**Figure 3:**
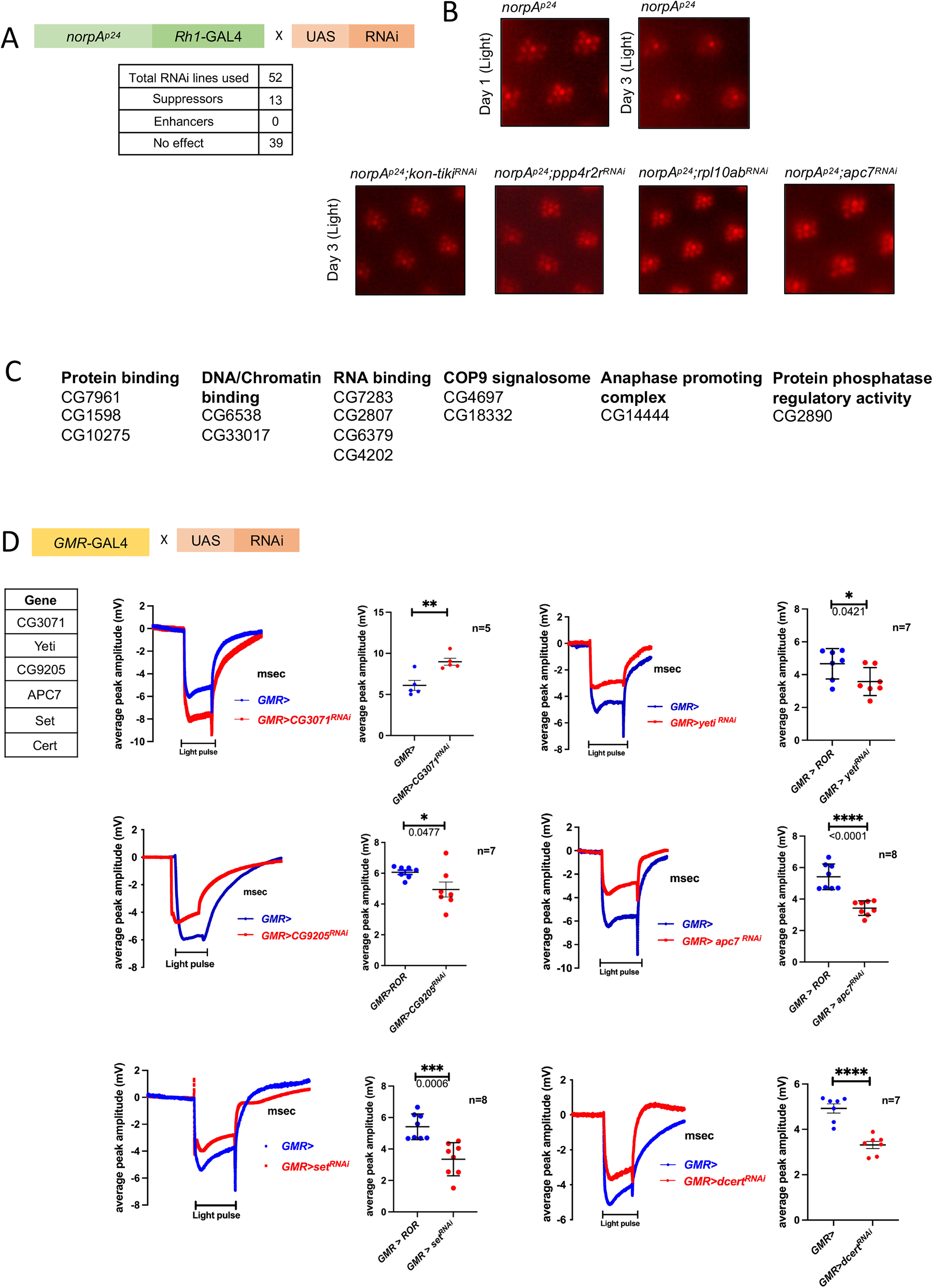
Genetic screen using *norpA^p24^*. (A) Scheme used to test for genetic interaction of each of the 52 *su(rdgB)* with *norpA^p24^* under illumination conditions (Constant light 2000 Lux). (B) *norpA^p24^* flies degenerate by day three under light conditions and examples of *su(RDGB)* candidates that suppressed *norpA^p24^* retinal degeneration phenotype. n=5 flies/RNAi line (C) Complete list 13 genes with their cellular functions that suppressed *norpA^p24^* phenotype. **ERG screen.** (D) Out of 52 candidates, five *su(RDGB)* showed reduced (*CG9205*, *Yeti*, *apc7*, *set*, *dcert*) and one (*CG3071*) showed higher ERG phenotype (traces and quantification shown) when downregulated in an otherwise wild type background. Number of flies used for the experimental set is mentioned along with the quantification. Scatter plots with mean ± SEM are shown. Statistical tests: Student’s unpaired t-test.

### ERG screen to identify *su(rdgB)* that may regulate phototransduction

A direct test of the role of a candidate in regulating phototransduction will be its ability, when depleted in an otherwise wild-type fly, to alter the electrical response to light. This can be monitored using electroretinograms (ERG) that are extracellular recordings that measure the electrical signal from the eye in response to a light stimulus (Vilinsky and Johnson, 2012). Any deviation of ERG amplitude when compared with that from a wild-type fly will imply that the interactor likely functions in the process of phototransduction. We downregulated each *su(rdgB)* using the eye specific promoter, GMR-GAL4 in an otherwise wild type background and measured ERG amplitudes. Out of 52 *su(rdgB),* GMR driven knockdown (in both of two independent RNAi lines) of five candidates (*CG9205*, *Yeti*, *APC7*, *Set*, *Cert*) showed a lower ERG amplitude and in one candidate (*CG3071*) a higher ERG amplitude compared to control flies (Figure 3D, Sup. Table S4). In the case of six additional *su(rdgB),* depletion with GMR-GAL4 resulted in a rough eye phenotype with the 1^st^ RNAi line (Sup. Fig1A i). When an 2^nd^ independent RNAi line was used, four (*Ars2*, *CG7483*, *cmtr1* and *secs*) out of six candidates showed lower ERG amplitude (Sup. Fig 1A iii). Rough eye phenotype after knocking down *Rpl10Ab* and *Sf3b1* with multiple RNAi lines point towards involvement of these genes in the eye development (Sup. Fig 1A ii).

### The spatial and temporal profile of *dcert* downregulation results in contrasting impact on *rdgB^9^* phenotypes

We previously noted that downregulation of *dcert* in *rdgB^9^* caused suppression of retinal degeneration when expressed using the Rh1 promoter (Figure 4A i, ii) although, the suppression in retinal degeneration was not sufficient to rescue ERG phenotype of *rdgB^9^* (Figure 4B i, ii). We retested this genetic interaction using a germline mutant allele of *dcert* (*dcert^1^*)(Rao et al., 2007)34. Surprisingly, the double mutant *rdgB^9^;dcert^1^* showed enhancement of retinal degeneration compared to *rdgB^9^* (Figure 4C i, ii). We tested these findings by using the same *dcert* RNAi line used in the screen (expressed using Rh1 Gal4) but this time with whole body expression of the RNAi using Actin-GAL4 which expresses throughout development beginning with embryogenesis. In *rdgB^9^;actin>dcert^RNAi^* we found enhancement of retinal degeneration such that by day 3 all photoreceptors except R7 were completely degenerated (Figure 4D i, ii). These findings suggest that *dcert* depletion more broadly in the fly across both space and time domains may have distinctive effects compared to a more restricted expression in post-mitotic adult photoreceptors using Rh1 GAL4.

**Figure 4:**
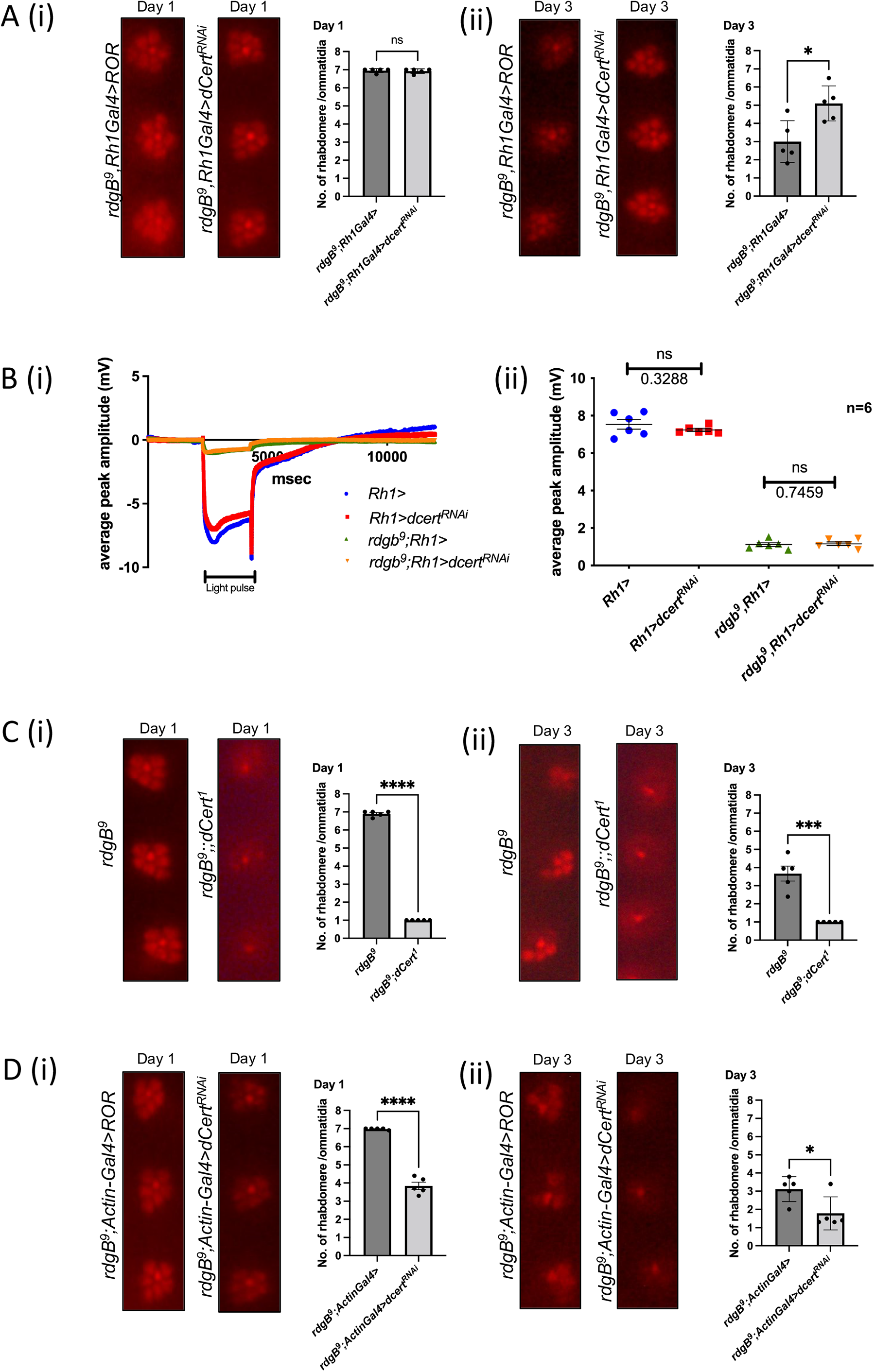
Spatial and temporal downregulation of *dcert* in *rdgB^9^*. (A) Suppression of retinal degeneration when RNAi lines were expressed using Rh1 enhancer. After eclosion flies were kept in the dark and assayed either on day one or day three (i) On day one there was no appreciable difference in two genotypes and rhabdomeres were intact (ii) on day three whereas control shows retinal degeneration downregulation of *dcert* in *rdgB^9^* suppressed the retinal degeneration observed in *rdgB^9^* control. (B) When subjected to ERG analysis, downregulation of *dcert* using *Rh1-GAL4* in the background of *rdgB^9^* did not suppress the ERG phenotype (i) ERG trace (ii) Quantification. n=6 flies, Scatter plots with mean ± SEM are shown. Statistical tests: Student’s unpaired t-test. (C) Double mutant of *rdgB^9^*;*dcert^1^* showed enhancement of retinal degeneration (i) (ii) By day one alone double mutant has severely enhanced retinal degeneration phenotype when compared to *rdgB^9^*. (D) Enhancement of retinal degeneration when *dcert* was downregulated with a whole-body Actin-Gal4 promoter in the *rdgB^9^* background (i) On day one rhabdomere loss is significant in the experimental files compared to control that worsens by day three and phenocopies the retinal degeneration present in the double mutant.

## Discussion

Neurodegeneration is a complex disease involving multiple layers of cellular and molecular process leading to the phenotype observed *in vivo*. Regardless of the part of the nervous system that is affected, be it the central or peripheral, conceptually, the processes leading to any neurodegeneration can be classified into two groups: (i) trigger steps-i.e those initial molecular or biochemical changes that initiate the process of degeneration (ii) Effector steps-i.e those steps that are subsequently part of the process that leads to loss of neuronal structure and consequently function. Identifying the molecular processes involved in each of these processes, is critical for developing strategies to manage neurodegenerative disorders. The *Drosophila* eye has been used in several settings for modelling neurodegenerations (Bonini and Fortini, 2003) such as those caused by repeat disorders such as Huntington’s disease and various ataxias, Alzheimer’s disease as well as primary degenerative disorders of the human retina (Xiong and Bellen, 2013). In the present study, we performed a genetic analysis to uncover the mechanisms of retinal degeneration underlying mutants in *rdgB*, that encodes a Class II PITP. Mutations in Class I PITP (PITPα) in mice result in a neurodegeneration phenotype (Hamilton et al., 1997) and recently human patients carrying mutations in VPS13 have been reported with neurodegenerative disorders (Ugur et al., 2020). Thus, the findings of our screen will inform on mechanisms of neurodegeneration.

To understand the cellular and molecular processes underlying retinal degeneration in *rdgB*, we depleted selected molecules using RNAi and scoring for suppression of the retinal degeneration. The candidates selected for screening were originally identified in a proteomic screen for interactors of VAP-A and VAP-B in cultured mammalian cells; however, the functional significance of their interaction with VAP was not known. Although previous studies have identified many VAP-interacting proteins in mammalian cell culture models by protein interaction studies, the functional relevance of these for *in vivo* function and neurodegeneration remains unknown. Using our *in vivo* analysis, we were able to identify a subset (52 out of 388) of these interactors in our proteomics screen that when depleted, suppressed the retinal degeneration in *rdgB^9^*. This finding underscores the value of an *in vivo* genetic screen in evaluating the functional effect of candidates identified *in vitro* to understanding the mechanisms of neurodegeneration. The human homologs in 13 of the *su(rdgB)* genes have previously been linked to human neurodevelopmental or neurodegenerative disorders (Table 1) and a large proportion of the 52 *su(rdgB)* have human homologs that show high expression in the human brain. Thus, the findings of this study could provide important insights into the mechanisms of human brain disorders.

Since our primary screen for suppressors of *rdgB* would identify molecules involved in both the trigger and effector steps of the degeneration process, it is essential to classify the identified suppressers into these two categories. Since *rdgB* mutants are known to effect photoreceptor physiology prior to the onset of retinal degeneration(Yadav et al., 2015), we reasoned that suppressors which work at the level of the trigger might also affect the electrical response to light, the physiological output of the photoreceptor. By this rationale, we found that 6 out of 52 suppressors when depleted in an otherwise wild-type background led to an altered electrical response to light; these suppressors are therefore likely to impact the processes by which RDGB functions in phototransduction. Examples of these include CG9205, Yeti, APC7, Set, Cert and CG3071. Two of these genes CG9205 (PH domain containing) and Cert (ceramide transfer protein) encode proteins with either ion binding or lipid transfer function and their ability to act as *su(rdgB)* may indicate a role for previous unidentified lipids and lipid transfer at MCS in phototransduction. By contrast Set (subunit of INHT complex that regulates histone acetylation), Yeti (a chromatin associated protein that interacts with the Tip60 chromatin remodelling complex) and CG3071 (snoRNA that positively regulates transcription by RNA polymerase 1) all likely exert their effect as *su(rdgB)* by modulating gene expression; some of the genes so regulated may impact phototransduction. A transcriptome analysis of *rdgB^9^* photoreceptors may help identify the relevant genes and the manner in which they regulate phototransduction.

To identify molecular mechanisms that regulate the effector steps of the degeneration process, we determined which of the *su(rdgB)* could also suppress another retinal degeneration mutant, norpAP*^24^*. Such *su(rdgB)* will likely represent molecules that participate in common effector steps of retinal degeneration shared by these two mutants. The 13 genes so identified represent several different functional classes. Prominent among these classes are RNA binding and DNA/chromatin binding proteins. Overall, a large percentage of *su(rdgB)* identified in our screen were of the class of RNA processing (CG1677, CG1542, CG7971, Cmtr1, Srrm234, Nop56, CG3071, Rpl10, Ars2, CG42458, SecS, CG9915, Sf3b1), RNA editing (Tailor, Sas10), RNA export (Phax), and RNA helicases (CG14443, CG7483). Interestingly, a role for RNA binding proteins such as ataxin-1 has been proposed in neuronal homeostasis and neurodegenerative processes and our finding may reflect a more general role for RNA binding/ homeostasis in neurodegenerative processes (Prashad and Gopal, 2021). A further large group of *su(rdgB)* belong to those regulating transcription (XNP, Fne, Yeti, TFIIFβ, TFIIEα, CG33017, Set, CG7839) and Sf3b1, Cmtr1, Rpl10Ab, TFIIFβ and Sas10 were among those candidates that additionally suppressed retinal degeneration in *norpA^p24^*. This finding suggests that regulated transcription may be important for maintaining neuronal homeostasis; this may be particularly significant since neurons are post-mitotic and transcriptional process and RNA turnover may collectively be key mechanisms for maintaining cellular homeostasis.

A third class of *su(rdgB)* were subunits of the COP9 signalosome (CSN1a, CSN2, CSN3 and CSN8 were identified in our screen). The COP9 signalosome acts as a signalling platform regulating cellular ubiquitylation status. The COP9 signalosome has been shown to play a key role in regulating *Drosophila* development through E3 ubiquitin ligases by deNEDDylation (Freilich et al., 1999). In addition, two E3 ubiquitin ligases family members were also identified in the genetic screen (i) APC7 which is a subunit of Anaphase promoting complex/Cyclosome that comprise of seven other subunits and is required to modulate cyclins levels during cell cycle (ii) CG32847, an uncharacterized gene belonging to the ‘Other RING domain ubiquitin ligases’ family of proteins. Ubiquitination could regulate the structure and function of proteins required for phototransduction; depletion of APC7 resulted in a reduction of the ERG amplitude supporting this mechanism. Alternatively, it is possible that ubiquitination regulated protein turnover may be part of the process of retinal degeneration. Interestingly, a key role for ubiquitination has been described in the context of neurodegeneration (Schmidt et al., 2021).

Overall, our screen uncovers a role for multiple molecular processes regulated by VAP interacting proteins that are required for maintaining lipid turnover and neuronal homeostasis in photoreceptors. It is important to note that our screen focused on VAP interacting proteins but there will also be non-VAP dependent processes that also contribute to lipid and neuronal homeostasis in photoreceptors. Alternative genetic screens will be required to map their role in photoreceptor maintenance. Collectively such studies will help advance our understanding of neurodegeneration in the context of lipid transfer protein function.

## Materials and Methods

### Protein pull-down and mass spectrometry analysis

Recombinant protein expression in *E. coli* and purification using plasmids encoding the MSP domain of VAP-A (8–212; WT and KD/MD mutant) and VAP-B (1–210; WT and KD/MD mutant) was previously described (di Mattia et al., 2020)18. For protein pull-down, the affinity resin was prepared by incubating 100 µg of recombinant protein with 20 µl of nickel beads (PureProteome Nickel magnetic beads, Merck) in 50 mM Pull-Down Buffer PDB (Tris–HCl pH 7.4, 50 mM NaCl, 1 mM EDTA, 1% Triton X-100, 5 mM imidazole, Complete protease inhibitor cocktail (Roche) and PhosSTOP (Roche)). The beads were then washed three times with the same buffer. 8 × 10^8^ HeLa cells were washed with 5 ml of TBS and lysed with 1 ml of PDB. After a 10-min incubation on ice, the protein extract was purified from cell debris by centrifugation (10 min; 9,500 g; 4°C). The protein extract was mixed with protein-coupled nickel beads and incubated for 2 h at 4°C under constant agitation. The beads were then washed three times with PDB, and proteins were eluted with Laemmli buffer. Proteins were precipitated with trichloroacetic acid and digested with Lys-C (Wako) and trypsin (Promega). The peptides were then analysed using an Ultimate 3000 nano-RSLC (Thermo Scientific) coupled in line with an Orbitrap ELITE (Thermo Scientific).

### SDS–PAGE, Western blot, and Coomassie blue staining

SDS–PAGE and Western blot analysis were performed as previously described (Alpy et al., 2005) using the following antibodies: rabbit anti-STARD3NL (1:1,000; pAbMENTHO-Ct-1545; (Alpy et al., 2001)43, rabbit anti-ORP1 (1:1,000; Abcam; ab131165), and mouse anti-actin (1:5,000; A1978 Merck). Coomassie blue staining was performed with PageBlue Protein Staining Solution (Thermo Fisher Scientific).

### *In silico* identification of potential conventional and Phospho FFAT motifs

The FFAT scoring algorithm used for Phospho-FFAT identification is based on the position weight matrix from Di Mattia, et.al (Di Mattia et al., 2020). For conventional FFAT sequences, the Phospho-FFAT matrix described in Di Mattia, et.al was modified in position 2 and 3 to assign a score of 4 to F and Y, and a score of 0 to D and E. These algorithms assign conventional and Phospho-FFAT scores to protein sequences. They are based on 19 continuous residues: six residues upstream, 7 residues forming the core and 6 residues downstream. An ideal sequence scores zero.

### Fly culture and stocks

Flies (*Drosophila melanogaster*) were reared on standard cornmeal, dextrose, yeast medium at 25°C and 50% relative humidity in a constant-temperature laboratory incubator. There was no internal illumination within the incubator and flies were subject to brief pulses of light only when the incubator doors were opened. To study light-dependent degeneration flies were exposed to light in an illuminated incubator at an intensity of 2000 lux. *rdgB^9^, P[w+,Rh1::GFP] ; Rh1-Gal4, UAS-Dicer2* and *norpA^p24^ ; Rh1-Gal4, UAS-Dicer2* were the strains used for the genetic screens.

### Fluorescent deep pseudopupil analysis

Pseudopupil analysis was carried out on flies after day 2 and day 4 post eclosion. Flies were immobilized using a stream of carbon dioxide and fluorescent pseudopupil analysis was carried out using an Olympus SZX12 stereomicroscope equipped with a fluorescent light source and green fluorescent protein (GFP) optics. Images were recorded using an Olympus digital camera.

### Optical neutralization

Flies were immobilized by cooling on ice. They were decapitated using a sharp razor blade and fixed on a glass slide using a drop of colourless nail varnish. The refractive index of the cornea was neutralized using a drop of immersion oil (*n*=1.516 at 23°C); images were observed using a 40× oil-immersion objective (Olympus, UPlanApo, 1.00 Iris) with antidromic illumination (Franceschini and Kirschfeld, 1971). Images were collected on an Olympus BX-41 upright microscope and recorded using an Olympus digital camera.

### Electroretinogram recordings

Flies were anesthetized and immobilized at the end of a disposable pipette tip using a drop of low melt wax. Recordings were done using glass microelectrodes filled with 0.8% w/v NaCl solution. Voltage changes were recorded between the surface of the eye and an electrode placed on the thorax. Following fixing and positioning, flies were dark adapted for 6 min. ERG was recorded with 1 second flashes of green light stimulus. Stimulating light was delivered from a LED light source within 5 mm of the fly’s eye through a fibre optic guide. Calibrated neutral density filters were used to vary the intensity of the light source. Voltage changes were amplified using a DAM50 amplifier (WPI) and recorded using pCLAMP 10.2. Analysis of traces was performed using Clampfit (Axon Laboratories).

## Acknowledgements

This work was supported by the Department of Atomic Energy, Government of India, under Project Identification No. RTI 4006, a Wellcome-DBT India Alliance Senior Fellowship to PR (IA/S/14/2/501540) and a Wellcome-DBT India Alliance Early Career Fellowship to SM (IA/E/17/1/503653). We thank the NCBS Imaging and Drosophila facilities for support. We thank Catherine Tomasetto and the other members of the Molecular and Cellular Biology of Breast Cancer team for helpful advice and discussions. We thank the IGBMC cell culture facility and proteomics platform (Luc Negroni, Frank Ruffenach and Bastien Morlet) for their excellent technical assistance. This work was supported by grants from the Agence Nationale de la Recherche ANR (grant ANR-19-CE44-0003; https://anr.fr/); This work of the Interdisciplinary Thematic Institute IMCBio, as part of the ITI 2021-2028 program of the University of Strasbourg, CNRS and Inserm, was supported by IdEx Unistra (ANR-10-IDEX-0002), and by SFRI-STRAT’US project (ANR 20-SFRI-0012) and EUR IMCBio (ANR-17-EURE-0023) under the framework of the French Investments for the Future Program.

---

**Supplementary Figure 1** (A) Second category of *su(rdgB)* was variable in either showing rough eye phenotype in 1^st^ RNAi line and ERG defects in the 2^nd^ independent RNAi line. (ii) *rpl10Ab* and *sf3b1* are the only candidates that consistently showed rough eye phenotype in two independent RNAi lines. (iii) ERG traces and quantifications of rest of the *su(rdgB)* with their respective RNAi line mentioned. Number of flies used for the experimental set is mentioned along with the quantification. Scatter plots with mean + SEM are shown. Statistical tests: Student’s unpaired t-test.

**Supplementary Table 1: List of VAP-A and VAP-B interacting proteins in an FFAT dependent manner**: Proteins identified by MS/MS after VAP-A and VAP-B pull-down. For each protein, the Uniprot ID, the name and the two best conventional FFAT and Phospho-FFAT scores are indicated. The position and the sequence of potential FFAT sequences are indicated. Proteins identified in VAP-A and VAP-B pull-down are labeled with a green background, and proteins identified in BioGRID 4.4.223 (Oughtred et al., 2019)as VAP partners are labeled in cyan. FFAT scores are color-coded with a scale from orange to blue (dark to light orange: 0-3, light to dark blue: 3.5->5). Acidic, phosphorylatable (S, T only), and aromatic (F, Y only) residues are shown in red, green and blue.

**Supplementary Table 2:** Total number of fly homologues/genes tested in the genetic screen. The genetic cross used to generate progeny for screening is shown at the top of the table. To perform this screen, we have used *rdgB^9^* recombined with Gal4 cassette under Rhodopsin 1 (Rh1) promoter at the 1^st^ chromosome (blue). This parental line was used to cross with each RNAi line expressing dsRNA against the specific fly gene (orange). Each fly gene is denoted with their specific CG number (www.flybase.org). Highlighted in red were those genotypes whose RNAi lines were not available. Highlighted in green were those genotypes where RNAi/genotype did not yield any flies after the cross.

**Supplementary Table 3:** Table showing each 52 *su(rdgB)* with their respective RNAi line tested for suppression in retinal degeneration in *norpA^p24^*. The genetic cross used to generate progeny for screening is shown at the top of the table. To perform this screen we have used *norpA^p24^* recombined with Gal4 cassette under Rhodopsin 1 (Rh1) promoter at the 1^st^ chromosome (green). This parental line was used to cross with each RNAi line expressing dsRNA against the specific fly gene (orange). Each fly gene is denoted with their specific CG number (www.flybase.org). KK/GD with a specific identifier number denotes the RNAi library generated by Vienna Drosophila Resource Centre (VDRC). Any suppression in retinal degeneration in *norpA^p24^* by downregulating the specific *su(rdgB) under* the Rh1 promoter is denoted by ‘Yes’.

**Supplementary Table 4:** Table showing each 52 *su(rdgB)* with their respective RNAi line tested for ERG/developmental phenotype when tested in an otherwise wild type background under GMR-Gal4. Each gene is denoted by their specific CG number. KK/GD denotes the RNAi library generated by Vienna Drosophila Resource Centre (VDRC) while TRiP lines denote the RNAi library procured from Bloomington Drosophila Resource Centre (BDSC). Where available Knockout (KO) lines were used. Phenotypes scored are denoted under ‘ERG’ column.

**Supplementary Table 5**: Table showing each 52 *su(rdgB)* with their potential FFAT motifs and their respective human homologue. For each human and fly protein, the Uniprot ID, the two best conventional and Phospho-FFAT scores are indicated. The position and the sequence of potential FFAT sequences are indicated. FFAT scores are color-coded with a scale from orange to blue (dark to light orange: 0-3, light to dark blue: 3.5->5). Acidic, phosphorylatable (S, T only), and aromatic (F, Y only) residues are shown in red, green and blue.

**Supplementary Table 6:** The levels of mRNA and protein of the identified genetics interactors of *rdgB* in brain. The data for mRNA expression and protein expression has been obtained from human protein atlas database (https://www.proteinatlas.org/)for the human homologues of the 52 genes reported as genetic interactors of *rdgB*. For the mRNA expression, the consensus TPM values from HPA in cerebral cortex and cerebellum (includes-HPA, gTEX and Fathom data) has been mentioned in column 4 and 5. The protein expression of the genes (mentioned as low, medium of high in HPA) has been shown for cerebral cortex and cerebellum in column 2 and 3. HPA reports the protein expression in various cell types of brain, however the region of the cerebral cortex and cerebellum with the highest expression has been used to report the protein expression. The cells marked in ‘yellow’ denote lack of data availability while cells marked in ‘blue’ denote low/no protein detected.

## References

Alpy, F., Stoeckel, M.-E., Dierich, A., Escola, J.-M., Wendling, C., Chenard, M.-P., Vanier, M. T., Gruenberg, J., Tomasetto, C. and Rio, M.-C. (2001). The Steroidogenic Acute Regulatory Protein Homolog MLN64, a Late Endosomal Cholesterol-binding Protein. Journal of Biological Chemistry 276, 4261–4269.

Alpy, F., Latchumanan, V. K., Kedinger, V., Janoshazi, A., Thiele, C., Wendling, C., Rio, M.-C. and Tomasetto, C. (2005). Functional Characterization of the MENTAL Domain. Journal of Biological Chemistry 280, 17945–17952.

Alpy, F., Rousseau, A., Schwab, Y., Legueux, F., Stoll, I., Wendling, C., Spiegelhalter, C., Kessler, P., Mathelin, C., Rio, M.-C., et al. (2013). STARD3/STARD3NL and VAP make a novel molecular tether between late endosomes and the ER. J Cell Sci. 10.1242/jcs.139295

Bonini, N. M. and Fortini, M. E. (2003). Human neurodegenerative disease modelling using Drosophila. Annu Rev Neurosci 26, 627–656.

Cabukusta, B., Berlin, I., van Elsland, D. M., Forkink, I., Spits, M., de Jong, A. W. M., Akkermans, J. J. L. L., Wijdeven, R. H. M., Janssen, G. M. C., van Veelen, P. A., et al. (2020). Human VAPome Analysis Reveals MOSPD1 and MOSPD3 as Membrane Contact Site Proteins Interacting with FFAT-Related FFNT Motifs. Cell Rep 33, 10.1016/j.celrep.2020.108475

Cockcroft, S. and Raghu, P. (2018). Phospholipid transport protein function at organelle contact sites. Curr Opin Cell Biol 53, 52–60.

Di Mattia, T., Martinet, A., Ikhlef, S., McEwen, A. G., Nominé, Y., Wendling, C., Poussin-Courmontagne, P., Voilquin, L., Eberling, P., Ruffenach, F., et al. (2020). FFAT motif phosphorylation controls formation and lipid transfer function of inter-organelle contacts. EMBO J 39, 10.15252/embj.2019104369

Dietzl, G., Chen, D., Schnorrer, F., Su, K.-C., Barinova, Y., Fellner, M., Gasser, B., Kinsey, K., Oppel, S., Scheiblauer, S., et al. (2007). A genome-wide transgenic RNAi library for conditional gene inactivation in Drosophila. Nature 448, 151–156.

Dudás, E. F., Huynen, M. A., Lesk, A. M. and Pastore, A. (2021). Invisible leashes: The tethering VAPs from infectious diseases to neurodegeneration. J Biol Chem 296, 10.1016/J.JBC.2021.100421

Fowler, P. C., Garcia-Pardo, M. E., Simpson, J. C. and O’Sullivan, N. C. (2019). NeurodegenERation: The Central Role for ER Contacts in Neuronal Function and Axonopathy, Lessons From Hereditary Spastic Paraplegias and Related Diseases. Front Neurosci 13, 10.3389/fnins.2019.01051

Franceschini, N. and Kirschfeld, K. (1971). Etude optique in vivo des éléments photorécepteurs dans l’œil composé de Drosophila. Kybernetik 8, 1–13.

Freilich, S., Oron, E., Kapp, Y., Nevo-Caspi, Y., Orgad, S., Segal, D. and Chamovitz, D. A. (1999). The COP9 signalosome is essential for development of Drosophila melanogaster. Current Biology 9, 1187–S4.

Georgiev, P., Garcia-Murillas, I., Ulahannan, D., Hardie, R. C. and Raghu, P. (2005). Functional INAD complexes are required to mediate degeneration in photoreceptors of the Drosophila rdgA mutant. J Cell Sci 118, 1373–1384.

Guillén-Samander, A. and De Camilli, P. (2022). Endoplasmic Reticulum Membrane Contact Sites, Lipid Transport, and Neurodegeneration. Cold Spring Harb Perspect Biol a041257. 10.1101/cshperspect.a041257

Hamilton, B. A., Smith, D. J., Mueller, K. L., Kerrebrock, A. W., Bronson, R. T., van Berkel, V., Daly, M. J., Kruglyak, L., Reeve, M. P., Nemhauser, J. L., et al. (1997). The vibrator Mutation Causes Neurodegeneration via Reduced Expression of PITPα: Positional Complementation Cloning and Extragenic Suppression. Neuron 18, 711–722.

Harayama, T. and Riezman, H. (2018). Understanding the diversity of membrane lipid composition. Nat Rev Mol Cell Biol 19, 281–296.

Hardie, R. and Raghu, P. (2001). Visual transduction in Drosophila. Nature. 413(6852) 186–93.

Harris, W. A. and Stark, W. S. (1977). Hereditary retinal degeneration in Drosophila melanogaster. A mutant defect associated with the phototransduction process. J.Gen.Physiol 69, 261–91.

Hotta, Y. and Benzer, S. (1970). Genetic dissection of the Drosophila nervous system by means of mosaics. Proc Natl Acad Sci USA 67, 1156–63.

Hu, Y., Flockhart, I., Vinayagam, A., Bergwitz, C., Berger, B., Perrimon, N. and Mohr, S. E. (2011). An integrative approach to ortholog prediction for disease-focused and other functional studies. BMC Bioinformatics 12, 357.

Kaiser, S. E., Brickner, J. H., Reilein, A. R., Fenn, T. D., Walter, P. and Brunger, A. T. (2005). Structural Basis of FFAT Motif-Mediated ER Targeting. Structure 13, 1035–1045.

Kim, Y. J., Guzman-Hernandez, M. L., Wisniewski, E. and Balla, T. (2015). Phosphatidylinositol-Phosphatidic Acid Exchange by Nir2 at ER-PM Contact Sites Maintains Phosphoinositide Signaling Competence. Dev Cell 33, 549–561.

Murphy, S. E. and Levine, T. P. (2016). VAP, a Versatile Access Point for the Endoplasmic Reticulum: Review and analysis of FFAT-like motifs in the VAPome. Biochimica et Biophysica Acta (BBA) – Molecular and Cell Biology of Lipids 1861, 952–961.

Oughtred, R., Stark, C., Breitkreutz, B. J., Rust, J., Boucher, L., Chang, C., Kolas, N., O’Donnell, L., Leung, G., McAdam, R., et al. (2019). The BioGRID interaction database: 2019 update. Nucleic Acids Res 47, D529–D541.

Pearn, M. T., Randall, L. L., Shortridge, R. D., Burg, M. G. and Pak, W. L. (1996). Molecular, Biochemical, and Electrophysiological Characterization of Drosophila norpA Mutants. Journal of Biological Chemistry 271, 4937–4945.

Peretti, D., Kim, S. H., Tufi, R. and Lev, S. (2020). Lipid Transfer Proteins and Membrane Contact Sites in Human Cancer. Front Cell Dev Biol 7.10.3389/fcell.2019.00371.

Perkins, L. A., Holderbaum, L., Tao, R., Hu, Y., Sopko, R., McCall, K., Yang-Zhou, D., Flockhart, I., Binari, R., Shim, H.-S., et al. (2015). The Transgenic RNAi Project at Harvard Medical School: Resources and Validation. Genetics 201, 843–852.

Prashad, S. and Gopal, P. P. (2021). RNA-binding proteins in neurological development and disease. RNA Biol 18, 972–987.

Prinz, W. A., Toulmay, A. and Balla, T. (2020). The functional universe of membrane contact sites. Nat Rev Mol Cell Biol 21, 7–24.

Raghu, P., Yadav, S. and Mallampati, N. B. N. (2012). Lipid signaling in Drosophila photoreceptors. Biochim Biophys Acta Mol Cell Biol Lipids 1821, 1154–1165.

Raghu, P., Basak, B. and Krishnan, H. (2021). Emerging perspectives on multidomain phosphatidylinositol transfer proteins. Biochim Biophys Acta Mol Cell Biol Lipids 1866.10.1016/j.bbalip.2021.158984

Rao, R. P., Yuan, C., Allegood, J. C., Rawat, S. S., Edwards, M. B., Wang, X., Merrill, A. H., Acharya, U. and Acharya, J. K. (2007). Ceramide transfer protein function is essential for normal oxidative stress response and lifespan. Proc. Natl. Acad. Sci. USA 104(27) 11364–9.

Rocha, N., Kuijl, C., van der Kant, R., Janssen, L., Houben, D., Janssen, H., Zwart, W. and Neefjes, J. (2009). Cholesterol sensor ORP1L contacts the ER protein VAP to control Rab7– RILP–p150Glued and late endosome positioning. Journal of Cell Biology 185, 1209–1225.

Schmidt, M. F., Gan, Z. Y., Komander, D. and Dewson, G. (2021). Ubiquitin signalling in neurodegeneration: mechanisms and therapeutic opportunities. Cell Death Differ 28, 570– 590.

Slee, J. A. and Levine, T. P. (2019). Systematic Prediction of FFAT Motifs Across Eukaryote Proteomes Identifies Nucleolar and Eisosome Proteins With the Predicted Capacity to Form Bridges to the Endoplasmic Reticulum. Contact 2, 251525641988313.

Stark, W. S., Chen, D.-M., Johnson, M. A. and Frayer, K. L. (1983). The *rdgB* Gene in Drosophila: retinal degeneration in different mutant alleles and inhibition of degeneration by *norpA*. J.Insect.Physiol 29(2): 123–131.

Ugur, B., Hancock-Cerutti, W., Leonzino, M. and De Camilli, P. (2020). Role of VPS13, a protein with similarity to ATG2, in physiology and disease. Curr Opin Genet Dev 65, 61–68.

Vihtelic, T. S., Hyde, D. R. and O’Tousa, J. E. (1991). Isolation and characterization of the Drosophila retinal degeneration B (rdgB) gene. Genetics 127, 761–768.

Vilinsky, I. and Johnson, K. G. (2012). Electroretinograms in *Drosophila*: A Robust and Genetically Accessible Electrophysiological System for the Undergraduate Laboratory. J Undergrad Neurosci Educ. 11(1):A149–57. Epub 2012 Oct 15.

Wilhelm, L. P., Wendling, C., Védie, B., Kobayashi, T., Chenard, M., Tomasetto, C., Drin, G. and Alpy, F. (2017). STARD3 mediates endoplasmic reticulum-to-endosome cholesterol transport at membrane contact sites. EMBO J 36, 1412–1433.

Wu, H., Carvalho, P. and Voeltz, G. K. (2018). Here, there, and everywhere: The importance of ER membrane contact sites. Science 361. Aug 3;361(6401):eaan5835.doi: 10.1126/science.aan5835.

Xiong, B. and Bellen, H. J. (2013). Rhodopsin homeostasis and retinal degeneration: lessons from the fly. Trends Neurosci 36, 652–660.

Yadav, S., Garner, K., Georgiev, P., Li, M., Gomez-Espinosa, E., Panda, A., Mathre, S., Okkenhaug, H., Cockcroft, S. and Raghu, P. (2015). RDGBα, a PtdIns-PtdOH transfer protein, regulates G-proteincoupled PtdIns(4,5)P_2_ signalling during *Drosophila* phototransduction. J Cell Sci 128, 3330–3344.

Yadav, S., Cockcroft, S. and Raghu, P. (2016). The Drosophila photoreceptor as a model system for studying signalling at membrane contact sites. Biochem Soc Trans 44, 447–451.

Yadav, S., Thakur, R., Georgiev, P., Deivasigamani, S., Krishnan, H., Ratnaparkhi, G. and Raghu, P. (2018). RDGBα localization and function at membrane contact sites is regulated by FFAT-VAP interactions. J Cell Sci 131. Jan 8;131(1):jcs207985.

