## supplemental file 1 for "A genetic screen to uncover molecular mechanisms underlying lipid transfer protein function at membrane contact sites and neurodegeneration"

A (i)

| Gene | 1st RNAi | Phenotype | 2nd RNAi | Phenotype |
|---|---|---|---|---|
| Ars2 | 106344/KK | Rough eye | 35204/TRiP | Less ERG |
| CG7483 | 108580/KK | Rough eye | 32444/TRiP | Less ERG |
| RpL10 | 109345/KK | Rough eye and necrosis | 23459/GD | Rough eye and necrosis |
| Sf3b1 | 110091/KK | Rough eye with heavy eye necrosis | 25162/GD | Rough eye with heavy eye necrosis |
| Cmtr1 | 103723/KK | Rough eye | 29611/GD | High ERG |
| SecS | 105727/KK | Rough eye | 38911/TRiP | Less ERG |

(ii)

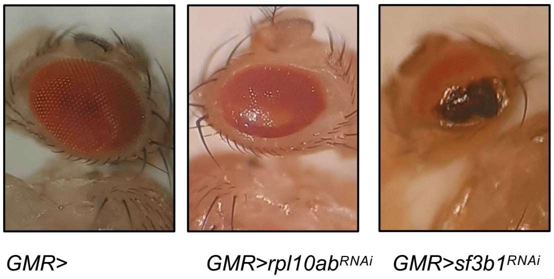


(iii)

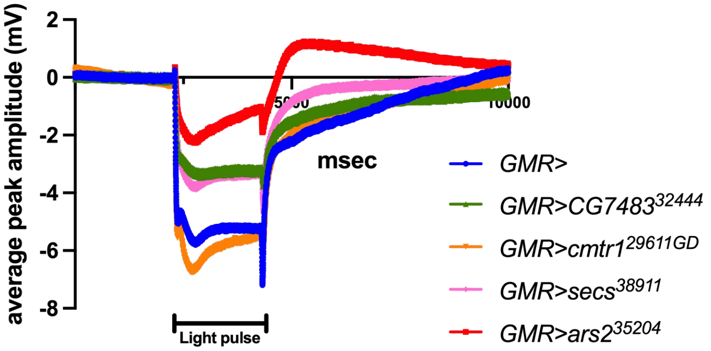


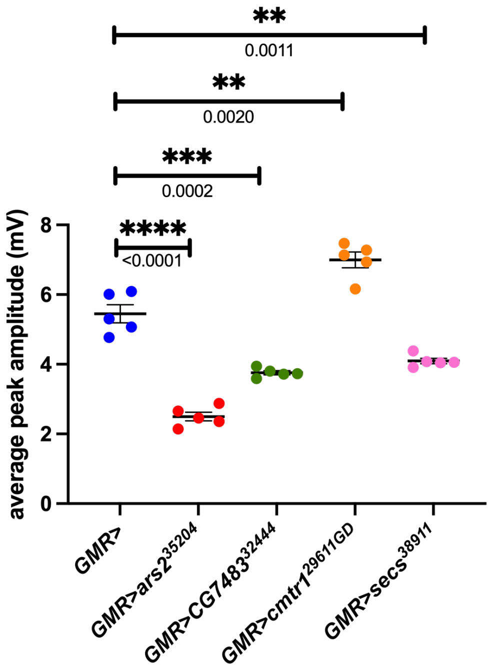


Sup. fig. 1
