## Supplemental Table S2 for "A genetic screen to uncover molecular mechanisms underlying lipid transfer protein function at membrane contact sites and neurodegeneration"

$rdgB^9$, Rh1-GFP | Rh1-GAL4 x UAS | RNAi

| | | | | | | | | | |
|---|---|---|---|---|---|---|---|---|---|
| 1 | CG3401 | 81 | CG8877 | 161 | CG9543 | 241 | CG11820 | 321 | CG4396 |
| 2 | CG9359 | 82 | CG32075 | 162 | CG1994 | 242 | CG3542 | 322 | CG6369 |
| 3 | CG1828 | 83 | CG33522 | 163 | CG3491 | 243 | CG4236 | 323 | CG8954 |
| 4 | CG2331 | 84 | CG10415 | 164 | CG44436 | 244 | CG5585 | 324 | CG16901 |
| 5 | CG13900 | 85 | CG9915 | 165 | CG1871 | 245 | CG15561 | 325 | CG3860 |
| 6 | CG2807 | 86 | CG4087 | 166 | CG2038 | 246 | CG7162 | 326 | CG2972 |
| 7 | CG7626 | 87 | CG17870 | 167 | CG17446 | 247 | CG3071 | 327 | CG17489 |
| 8 | CG10417 | 88 | CG31196 | 168 | CG3347 | 248 | CG4488 | 328 | CG3458 |
| 9 | CG16742 | 89 | CG9556 | 169 | CG17440 | 249 | CG8176 | 329 | CG7516 |
| 10 | CG8274 | 90 | CG2173 | 170 | CG1542 | 250 | CG4364 | 330 | CG43665 |
| 11 | CG3312 | 91 | CG1582 | 171 | CG11444 | 251 | CG44086 | 331 | CG8414 |
| 12 | CG6049 | 92 | CG9323 | 172 | CG4438 | 252 | CG4049 | 332 | CG18304 |
| 13 | CG4299 | 93 | CG32217 | 173 | CG7686 | 253 | CG6223 | 333 | CG30170 |
| 14 | CG5784 | 94 | CG17437 | 174 | CG8545 | 254 | CG7538 | 334 | CG8915 |
| 15 | CG17593 | 95 | CG10931 | 175 | CG1276 | 255 | CG7108 | 335 | CG4379 |
| 16 | CG3732 | 96 | CG9594 | 176 | CG10254 | 256 | CG3335 | 336 | CG10540 |
| 17 | CG6227 | 97 | CG8103 | 177 | CG8635 | 257 | CG11130 | 337 | CG4878 |
| 18 | CG2503 | 98 | CG14884 | 178 | CG11266 | 258 | CG40494 | 338 | CG5519 |
| 19 | CG6708 | 99 | CG9423 | 179 | CG17768 | 259 | CG42522 | 339 | CG32707 |
| 20 | CG1406 | 100 | CG9205 | 180 | CG6932 | 260 | CG5642 | 340 | CG6964 |
| 21 | CG14472 | 101 | CG17249 | 181 | CG8863 | 261 | CG31864 | 341 | CG1430 |
| 22 | CG34133 | 102 | CG7061 | 182 | CG12020 | 262 | CG12264 | 342 | CG33967 |
| 23 | CG4548 | 103 | CG6724 | 183 | CG9828 | 263 | CG15224 | 343 | CG6701 |
| 24 | CG10840 | 104 | CG7597 | 184 | CG11920 | 264 | CG32505 | 344 | CG6967 |
| 25 | CG16941 | 105 | CG33556 | 185 | CG13667 | 265 | CG3183 | 345 | CG6379 |
| 26 | CG4799 | 106 | CG2890 | 186 | CG13298 | 266 | CG5904 | 346 | CG13096 |
| 27 | CG10986 | 107 | CG7843 | 187 | CG4119 | 267 | CG3029 | 347 | CG5728 |
| 28 | CG7706 | 108 | CG1433 | 188 | CG8161 | 268 | CG16792 | 348 | CG8781 |
| 29 | CG2469 | 109 | CG10887 | 189 | CG7207 | 269 | CG9344 | 349 | CG3949 |
| 30 | CG2925 | 110 | CG10418 | 190 | CG6759 | 270 | CG7917 | 350 | CG5271 |
| 31 | CG8548 | 111 | CG4528 | 191 | CG10811 | 271 | CG9397 | 351 | CG42341 |
| 32 | CG12225 | 112 | CG11418 | 192 | CG10192 | 272 | CG33106 | 352 | CG4279 |
| 33 | CG8892 | 113 | CG1091 | 193 | CG12372 | 273 | CG6904 | 353 | CG8069 |
| 34 | CG12272 | 114 | CG7163 | 194 | CG5403 | 274 | CG2257 | 354 | CG11738 |
| 35 | CG10080 | 115 | CG8817 | 195 | CG7467 | 275 | CG14739 | 355 | CG2253 |
| 36 | CG5216 | 116 | CG3889 | 196 | CG9537 | 276 | CG30342 | 356 | CG16788 |
| 37 | CG11990 | 117 | CG5627 | 197 | CG5800 | 277 | CG2790 | 357 | CG34334 |
| 38 | CG7028 | 118 | CG8427 | 198 | CG2656 | 278 | CG9124 | 358 | CG8282 |
| 39 | CG8400 | 119 | CG1017 | 199 | CG10222 | 279 | CG10890 | 359 | CG6905 |
| 40 | CG1513 | 120 | CG4697 | 200 | CG9548 | 280 | CG6315 | 360 | CG1616 |
| 41 | CG8725 | 121 | CG8956 | 201 | CG6554 | 281 | CG4039 | 361 | CG10370 |
| 42 | CG7769 | 122 | CG11654 | 202 | CG6677 | 282 | CG7006 | 362 | CG11228 |
| 43 | CG6222 | 123 | CG9977 | 203 | CG4806 | 283 | CG6946 | 363 | CG4063 |
| 44 | CG31935 | 124 | CG8625 | 204 | CG1575 | 284 | CG5824 | 364 | CG4913 |
| 45 | CG9484 | 125 | CG7727 | 205 | CG4886 | 285 | CG5605 | 365 | CG9635 |
| 46 | CG1249 | 126 | CG10161 | 206 | CG1866 | 286 | CG4817 | 366 | CG5670 |
| 47 | CG7728 | 127 | CG4810 | 207 | CG9916 | 287 | CG3605 | 367 | CG2747 |
| 48 | CG5504 | 128 | CG1815 | 208 | CG8336 | 288 | CG5654 | 368 | CG11238 |
| 49 | CG3909 | 129 | CG5033 | 209 | CG7768 | 289 | CG5208 | 369 | CG32146 |
| 50 | CG13097 | 130 | CG4202 | 210 | CG17266 | 290 | CG5229 | 370 | CG3198 |
| 51 | CG17255 | 131 | CG10281 | 211 | CG5258 | 291 | CG3025 | 371 | CG7971 |
| 52 | CG8583 | 132 | CG6538 | 212 | CG31301 | 292 | CG11290 | 372 | CG5444 |
| 53 | CG7757 | 133 | CG30349 | 213 | CG12608 | 293 | CG1894 | 373 | CG7015 |
| 54 | CG10754 | 134 | CG14444 | 214 | CG9123 | 294 | CG1691 | 374 | CG13472 |
| 55 | CG14894 | 135 | CG1559 | 215 | CG14805 | 295 | CG7961 | 375 | CG4539 |
| 56 | CG3035 | 136 | CG5786 | 216 | CG2508 | 296 | CG34126 | 376 | CG11727 |
| 57 | CG42665 | 137 | CG33208 | 217 | CG31687 | 297 | CG7283 | 377 | CG13185 |
| 58 | CG8421 | 138 | CG7035 | 218 | CG3733 | 298 | CG3843 | 378 | CG32409 |
| 59 | CG9351 | 139 | CG7907 | 219 | CG9253 | 299 | CG4581 | 379 | CG32211 |
| 60 | CG8939 | 140 | CG1554 | 220 | CG42458 | 300 | CG4622 | 380 | CG10390 |
| 61 | CG5659 | 141 | CG4152 | 221 | CG8092 | 301 | CG9346 | 381 | CG16973 |
| 62 | CG4170 | 142 | CG43658 | 222 | CG10850 | 302 | CG11271 | 382 | CG2380 |
| 63 | CG1677 | 143 | CG32251 | 223 | CG7041 | 303 | CG1527 | 383 | CG34401 |
| 64 | CG40218 | 144 | CG1427 | 224 | CG15636 | 304 | CG7036 | 384 | CG3522 |
| 65 | CG11427 | 145 | CG2238 | 225 | CG8120 | 305 | CG33526 | 385 | CG5589 |
| 66 | CG9198 | 146 | CG4849 | 226 | CG8409 | 306 | CG13849 | 386 | CG1507 |
| 67 | CG18332 | 147 | CG1598 | 227 | CG6990 | 307 | CG10206 | 387 | CG10528 |
| 68 | CG8571 | 148 | CG11985 | 228 | CG10275 | 308 | CG9018 | 388 | CG4510 |
| 69 | CG40351 | 149 | CG11188 | 229 | CG6349 | 309 | CG31022 | 389 | CG3725 |
| 70 | CG33217 | 150 | CG8610 | 230 | CG4954 | 310 | CG4164 | 390 | CG7946 |
| 71 | CG10223 | 151 | CG7483 | 231 | CG11417 | 311 | CG6833 | 391 | CG5434 |
| 72 | CG5205 | 152 | CG9075 | 232 | CG2028 | 312 | CG2791 | 392 | CG17603 |
| 73 | CG2093 | 153 | CG7421 | 233 | CG2048 | 313 | CG9805 | 393 | CG9062 |
| 74 | CG3696 | 154 | CG3817 | 234 | CG9962 | 314 | CG12792 | | |
| 75 | CG6876 | 155 | CG10955 | 235 | CG7094 | 315 | CG5004 | | |
| 76 | CG5899 | 156 | CG12498 | 236 | CG2577 | 316 | CG9983 | | |
| 77 | CG32346 | 157 | CG31703 | 237 | CG12147 | 317 | CG6354 | | |
| 78 | CG7839 | 158 | CG31702 | 238 | CG2685 | 318 | CG12749 | | |
| 79 | CG1528 | 159 | CG5794 | 239 | CG7263 | 319 | CG4262 | | |
| 80 | CG14443 | 160 | CG3730 | 240 | CG6699 | 320 | CG3151 | | |

No available RNAi lines

Genotypes where no larvae/fly emerged after the cross

Table S2
