## Supplemental Table S4 for "A genetic screen to uncover molecular mechanisms underlying lipid transfer protein function at membrane contact sites and neurodegeneration"

*GMR*-GAL4 x UAS RNAi

| Gene | RNAi | ERG | 2ND RNAi/KO | ERG |
|---|---|---|---|---|
| CG8069 | 28189/GD | No | | |
| CG4548 | 101568/KK | No | | |
| CG7961 | 35305/GD | No | | |
| CG7843 | 106344/KK | **Rough eye** | 35204/TRiP | **Less** |
| CG42665 | 106344/KK | No | | |
| CG8092 | 28196/GD | No | | |
| CG42458 | 106608/KK | No | | |
| CG42522 | 33370/TRiP | No | | |
| CG1677 | 109697/KK | No | | |
| CG14443 | 105254/KK | No | | |
| CG1542 | 104575/KK | No | | |
| CG9825 | 105868/KK | No | | |
| CG9205 | 29079/GD | **Less** | *CG9205*$^{KO}$ | **Less** |
| CG7971 | 101384/KK | No | | |
| CG4799 | 102627/KK | No | | |
| CG9915 | 103731/KK | No | | |
| CG13849 | 103738/KK | **High** | 51775/GD | **No** |
| CG9181 | 108888/KK | No | | |
| CG4396 | 101508/KK | **Rough eye** | 28784/TRiP | **No** |
| CG33967 | 106507/KK | No | | |
| CG13176 | 39769/GD | No | | |
| CG3071 | 107206/KK | **High** | No RNAi line | |
| CG1598 | 110555/KK | No | | |
| CG40218 | 102960/KK | **Less** | No RNAi line | |
| CG4697 | 34308/GD | No | | |
| CG14444 | 110729/KK | **Less** | 17622/GD | **Less** |
| CG2890 | 105399/KK | **Rough eye** | 26296/TRiP | **No** |
| CG7283 | 109345/KK | **Rough eye** | 23459/GD | **Rough eye** |
| CG2807 | 110091/KK | **Rough eye** | 25162/GD | **Rough eye** |
| CG6538 | 12602/GD | No | | |

| Gene | RNAi | ERG | 2ND RNAi/KO | ERG |
|---|---|---|---|---|
| CG18332 | 101516/KK | No | | |
| CG6379 | 103723/KK | **Rough eye** | 29611/GD | **High** |
| CG1427 | 105727/KK | **Rough eye** | 38911/TRiP | **Less** |
| CG10275 | 106680/KK | No | | |
| CG2790 | 101619/KK | No | | |
| CG10415 | 100572/KK | No | | |
| CG11228 | 104169/KK | No | | |
| CG1091 | 107175/KK | No | | |
| CG33106 | 103411/KK | No | | |
| CG33208 | 105837/KK | No | | |
| CG15224 | 106845/KK | No | | |
| CG17593 | 106469/KK | No | | |
| CG33017 | 103968/KK | No | | |
| CG4299 | 108987/KK | **Less** | 77433/TRiP | **Less** |
| CG7207 | 35579/TRiP | **Less** | *dcert*$^{1}$ | **Less** |
| CG4202 | 103352/KK | No | | |
| CG9977 | 106749/KK | No | | |
| CG32847 | 104294/KK | No | | |
| CG7839 | 105979/KK | No | | |
| CG7483 | 108580/KK | **Rough eye** | 32444/TRiP | **Less** |
| CG9556 | 48044/GD | No | | |
| CG3825 | 107545/KK | No | | |

Table S4
